# MET Inhibitor Capmatinib Radiosensitizes MET Exon 14-Mutated and MET-Amplified Non-Small Cell Lung Cancer

**DOI:** 10.1101/2023.10.26.564232

**Authors:** Shrey Ramesh, Ahmet Cifci, Saahil Javeri, Rachel Minne, Colin A. Longhurst, Kwangok P. Nickel, Randall J. Kimple, Andrew M. Baschnagel

**Author notes:** **Corresponding Author Names & Email Addresses:** Randall J. Kimple, MD, PhD, Department of Human Oncology, University of Wisconsin-Madison School of Medicine and Public Health, 600 Highland Avenue, K4/B100-0600, Madison, WI 53792,;Andrew M. Baschnagel, MD, Department of Human Oncology, University of Wisconsin-Madison School of Medicine and Public Health, 600 Highland Avenue, K4/B100-0600, Madison, WI 53792. Author Responsible for Statistical Analysis Name & Email Address: Colin A. Longhurst, BS. Shrey Ramesh and Ahmet Cifci contributed equally as co-first authors.

## Abstract

**Purpose:** The objective of this study was to investigate the effects of inhibiting the MET receptor with capmatinib, a potent and clinically relevant ATP-competitive tyrosine kinase inhibitor, in combination with radiation in MET exon 14-mutated and MET-amplified non-small cell lung (NSCLC) cancer models.

**Methods and Materials:** *In vitro* effects of capmatinib and radiation on cell proliferation, colony formation, MET signaling, apoptosis, and DNA damage repair were evaluated. *In vivo* tumor responses were assessed in cell line xenograft and patient-derived xenograft models. Immunohistochemistry (IHC) was used to confirm *in vitro* results.

**Results:** *In vitro* clonogenic survival assays demonstrated radiosensitization with capmatinib in both MET exon 14-mutated and MET-amplified NSCLC cell lines. No radiation-enhancing effect was observed in MET wild-type NSCLC and human bronchial epithelial cell line. Minimal apoptosis was detected with the combination of capmatinib and radiation. Capmatinib plus radiation compared to radiation alone resulted in inhibition of DNA double-strand break repair as measured by prolonged expression of γH2AX. *In vivo*, the combination of capmatinib and radiation significantly delayed tumor growth compared to vehicle control, capmatinib alone, or radiation alone. IHC indicated inhibition of phospho-MET and phospho-S6 and a decrease in Ki67 with inhibition of MET.

**Conclusions:** Inhibition of MET with capmatinib enhanced the effect of radiation in both MET exon 14-mutated and MET-amplified NSCLC models.

## Introduction

MET is a proto-oncogene encoding a receptor tyrosine kinase that plays a critical role in cell proliferation, migration, survival, and differentiation. The MET protein is activated by its ligand, hepatocyte growth factor (HGF), which leads to the activation of downstream signaling pathways, including the PI3K/AKT and MAPK/ERK pathways. MET dysregulation frequently occurs in many cancers leading to tumor progression. In patients with non-small cell lung cancer (NSCLC), MET is mutated in 3-4%(1–3) and amplified in 3-6%(4–7) of tumors. The most common MET mutation occurs in exon 14, which leads to impaired degradation of the protein resulting in over-expression and activation of the receptor. Both MET exon 14 skipping mutations and MET amplification can be successfully inhibited with newly FDA-approved small molecule inhibitors.

Several selective MET tyrosine kinase inhibitors have shown efficacy in patients with MET-altered NSCLC.(8–11) Both capmatinib and tepotinib are now FDA approved as first-line therapy in patients with metastatic NSCLC harboring MET exon 14 skipping mutations.(8, 9) Additionally, both agents show activity in patients with NSCLC with high MET amplification.

Combination therapy with MET inhibitors and other targeted therapies, such as epidermal growth factor receptor (EGFR) inhibitors, are also being investigated as potential treatment options for metastatic NSCLC patients with co-occurring alterations and following the inevitable development of acquired resistance to current small molecule inhibitors.(12–15)

Multiple preclinical studies have now demonstrated that inhibition of MET signaling can increase the sensitivity of cancer cells to radiation in various tumor types.(16–26) Two studies in NSCLC have shown *in vitro* radiosensitization with two different selective small-molecule inhibitors of MET. Li et al. have shown that AMG-458 radiosensitizes one MET expressing cell line (NCI-H441) (17) and Bhardwaj et al. have shown that MK-8033 radiosensitizes two MET-amplified cell lines.(16) The limitations of these studies are that both drugs evaluated are not available in patients and neither of these two studies investigated the effect in MET exon 14 models or performed *in vivo* experiments. We have previously shown *in vivo* radiation enhancement with savolitinib (26), but to the best of our knowledge, no preclinical or clinical studies have examined the potential synergistic effects of the first-line MET inhibitor, capmatinib, in combination with radiation therapy. Combining radiation with a MET inihibitor may be a useful strategy in both the locally advanced and metastatic setting.

The objective of this study was to evaluate the radiation enhancement response of combining the clinically relevant MET inhibitor capmatinib with radiation in both MET exon 14 and MET-amplified NSCLC cell lines and patient-derived xenograft (PDX) models. We further tested whether capmatinib and radiation combination influences MET signaling and investigated the mechanism of cell death by examining DNA damage repair and apoptotic pathways.

## Methods and Materials

### Cell lines, patient-derived xenografts, and drug

Three human NSCLC cell lines, one human non-tumorigenic lung epithelial cell line, and two NSCLC PDXs were used in this study. A549, derived from lung adenocarcinoma, has a KRAS mutation, is MET wild type, and was obtained from ATCC (Manassas, VA). EBC-1 was derived from squamous cell carcinoma, has high MET amplification(27), and was obtained from JCRB Cell Bank (Xenotech, Kansas City, KS). Beas-2B was derived from non-tumorigenic human lung epithelium, is MET wild type and obtained from ATCC (Manassas, VA). The UW-lung-21cell line and PDX were derived from a human lung adenocarcinoma brain metastasis harboring a MET exon 14 skipping mutation(26). For development of the UW-lung-21cell line, tumors taken from early PDX passages were dissociated into single cell suspensions and seeded in 6-well plates containing media (RPMI with 10% fetal bovine serum, 1% L-glutamine, 1% penicillin/streptomycin, and 2.5 μg/mL amphotericin B) and incubated at 37 °C in a humidified atmosphere of 5% CO_2_. Media was changed regularly, and cells were passaged once they reached 70% confluence. Mycoplasma testing and short tandem repeat profiling were performed routinely. BM016-16 PDX was derived from a NSCLC brain metastasis tumor with MET amplification (FiSH 11.7 MET/CEP7) and was generously provided by Dr. Timothy F. Burns and Dr. Laura Stabile (University of Pittsburgh). All cell lines had their identity confirmed via short tandem repeat profiling analysis (Supplemental Table 1), were maintained, used mycoplasma-free media at a lower passage, and were cultured as described in Supplemental Table 2. Capmatinib was purchased from Selleck Chemicals (Houston, TX).

### Irradiation

Cells were irradiated using a Xstrahl X-ray System, Model RS225 (Xstrahl, UK) at a dose rate of 3.27 Gy/min at 30 cm FSD, tube voltage of 195 kV, current of 10 mA, and filtration with 3 mm Al at doses ranging from 2-6 Gy. For the xenograft experiments, animals were irradiated with a Xstrahl X-ray System, Model CIX3 (Xstrahl, UK). Each 2 Gy fraction was delivered at a 2.04 Gy/min with 300 kV and 10 mA at 40 cm FSD with 1 mm Cu filter. The delivered dose rate was confirmed monthly by ionization chamber by the University of Wisconsin Medical Radiation Research Center Research Center Calibration lab. Mice were shielded with custom-built lead jigs to isolate exposure to the rear quarter of the body.

### Cell proliferation

Cells were plated in 96-well plates at densities ranging from 3,000 to 6,000 cells per well according to each cell growth rate. Twenty-four h after-plating, cells were treated with indicated doses of capmatinib or control (DMSO) and incubated for 72 h. Afterwards, Cell Counting Kit-8 reagent was added (Dojindo Molecular Technologies) and absorbance measured at 450 nm on a SpectraMax i3 plate reader (Molecular Devices). The absorbance of treated wells was normalized to control wells and the half-maximal inhibitory concentration (IC50) values were calculated.

### Western blot analysis

Cells were seeded in 10-cm dishes and treated with either control (DMSO) or 5 nM capmatinib after overnight incubation. Irradiation (2 Gy) was applied 1 h after control or capmatinib treatment, and cells were harvested 4 or 24 h after radiation or mock treatment. Cell lysates were generated as previously described(28, 29). The specific antibodies and sources are listed in Supplemental Table 3.

### Clonogenic survival assay

Cells were seeded into 6-well plates at specific densities, incubated overnight, and then irradiated as indicated after 1 h of capmatinib or control treatment. The next day, media was removed and replaced with fresh, drug-free media. Once colonies averaged 50 or more cells (12–20 days) in the control wells, plates were fixed and stained with 1% (w/v) crystal violet in methanol, imaged, and colonies of 50 or more cells were counted. Survival curves were generated after normalizing for capmatinib-induced cell death. The clonogenic survival curve for each condition was fitted to a linear quadratic model [Y = e - (A × X + B × X^2^)] according to a least-squares fit, weighted to minimize the relative distances squared. Each point represents the mean surviving fraction calculated from three replicates and the assays have been repeated at least three times. Radiation dose enhancement factor (DEF) was calculated at 10% survival levels by dividing the mean radiation dose for control conditions by the mean radiation dose after drug exposure. A value >1.0 indicates enhancement of radiosensitivity.

### **γ**H2AX assay

γH2AX immunofluorescence assays were performed as described previously(29). Briefly, cells were plated on coverslips in 24-well plates, treated with 10 nM capmatinib, and irradiated with 4 Gy 1 h after the drug treatment the next day. Then cells were fixed at 1, 4, or 24 h after the treatment, permeabilized, and probed with primary and secondary antibodies. The specific antibodies and sources are listed in Supplemental Table 3. Slides were imaged at 60X magnification using a Leica SP8 confocal WLL STED microscope. γH2AX foci per cell were counted using FIJI(30) with at least 150 cells from three replicates. The intensity threshold was set in the red channel using the control group for each experiment. A binary image was created using the threshold and foci were counted automatically excluding particles below a minimum size threshold of 0.0002 micron.

### Comet assay

Cells were plated in 6-well plates at 100,000 cells/well, incubated overnight, and then treated with either control or 10 nM capmatinib, 4 Gy of irradiation, or 10 nM of capmatinib followed 1 hr by 4 Gy irradiation and collected a 4 or 24 hr later. CometAssay® ES II electrophoresis system (R&D systems) was used as previously described (31). Cells were trypsinized, counted, embedded in LMAgarose on two well slides at a density of 500 cells/well, and lysed overnight at 4°C. Cells were immersed in 1X neutral electrophoresis buffer at 4°C for 30 minutes, and electrophoresis was then performed at 21 V for 45 minutes. Cells were immersed in DNA precipitation solution for 30 minutes at room temperature, followed by immersion in 70% ethanol for 30 minutes at room temperature. Samples were dried at 37°C for 15 minutes and each sample was stained using 100μL of diluted SYBR® Gold for 30 minutes in the dark at room temperature. Cells were allowed to dry completely at 37°C before image analysis. Images of cells were taken on a Nikon Intensilight Fluorescence Microscope at 20X. At least 50 cells in each treatment group at each timepoint were imaged and analyzed. The olive tail moment was calculated as: Tail DNA % ×Tail Moment Length, using the Comet Score 2.0 software. The Tail DNA % is the intensity of the tail DNA and the Tail Moment Length is the distance between the center of the comet head and the center of the comet tail. Three biological samples with a total of over 150 cells were counted and used to calculate average olive tail moment values.

### Apoptosis

Apoptosis was detected using Annexin V/FITC staining kit (Thermo Fisher). Briefly, cells were seeded in 6-well plates, incubated overnight, treated with 10 nM capmatinib, and irradiated with 4 Gy 1 h after drug treatment. Both early and late apoptosis was detected 48 h after treatment according to the manufacturer’s instructions using an Attune Next Flow Cytometer (Thermo Fisher). Flow cytometry data were analyzed using FlowJo.

### Xenograft growth delay studies

Six- to eight-week-old female Hsd:athymic Nude-Foxn1nu mice (Envigo) were used for growth delay studies. Mice were kept in the Association for Assessment and Accreditation of Laboratory Animal Care-approved Wisconsin Institute for Medical Research Animal Care Facility. All animal studies were conducted in accordance with, and with the approval of, the University of Wisconsin Institutional Animal Care and Use Committee (IACUC). Animals were housed in specific pathogen-free rooms, and their clinical health was evaluated weekly. Studies involving the mice were carried out in accordance with an animal protocol approved by the University of Wisconsin.

Tumor cells (EBC-1) or PDX tissue slurry (UW-lung-21and BM016-16) mixed with Matrigel (1:1; Corning) were injected subcutaneously into the bilateral hind flanks of athymic nude mice. When tumors grew to a mean volume of 150-300 mm^3^, mice were randomized into control, radiation alone (2 Gy per fractions for 5 days), capmatinib (20 mg/kg for 5 days), or radiation plus capmatinib. Capmatinib was administered by oral gavage. Tumor volume was measured twice weekly with digital vernier caliper and calculated using V = (π/6) X (large diameter) X (small diameter) ^two^. Each experimental group contained 8 to 10 mice.

### Immunohistochemistry

After harvesting, tumor was fixed in 10% neutral-buffered formalin and embedded in paraffin blocks. 5-μm sections of formalin-fixed, paraffin-embedded samples were deparaffinized with Xylene and hydrated through graded solutions of ethanol. Antigen retrieval was conducted in sodium citrate retrieval buffer (pH 6.0) followed by washing in running water. The slides were washed in PBS and then incubated with a 0.3% hydrogen peroxide solution. Blocking was carried out using 10% goat serum in PBS and then incubated with the primary antibody (Supplemental Table 3) diluted in 1% goat serum in PBS containing 0.1% Triton X-100 overnight at 4°C. The slides were washed with PBS the next day; secondary antibody (Signal Stain Boost IHC Detection Reagent (HRP, Rabbit) CST #8114) was used. Staining was detected using diaminobenzidine (Vector Laboratories, Inc. #SK-4100). The slides were counterstained with 1:10 hematoxylin (Thermo Scientific, #TA-125-MH) solution for 2 min, then dehydrated in ethanol and xylene solutions and sections were covered with coverslip with Cytoseal (Thermo Scientific #8312-4). Details of the antibodies used are listed in Supplemental Table 3. Images were obtained on an Olympus BX51 microscope (Olympus America, Inc.).

Quantification of Ki67 was completed using FIJI to identify areas of positive staining(30).

Images underwent color deconvolution to isolate DAB staining, and the threshold tool was applied to automatically calculate the percentage area positively stained. The threshold was set with the control group for each experiment.

### Statistical analyses

*In vitro* experiments were repeated three times and statistical analyses were carried out using a Student’s t-test or one-way ANOVA. Differences in radiation clonogenic survival curves were determined using the extra sum-of-squares F test. Data are presented as the mean ± standard error of the mean (SEM). A *P* value of < .05 was considered statistically significant. All graphs and *in vitro* analyses were performed and graphed using GraphPad Prism version 9.0 (GraphPad Software). For the longitudinal xenograft tumor growth volume assessment, generalized linear mixed models (with a log-link function) were fit to the data using the “lme4” package in R (V 4.0.3). The fixed-effect portion of the models was parameterized such that all main and interactive effects between treatment groups (vehicle control, capmatinib, radiation, and capmatinib plus radiation) and time were estimated, whereas the random-effect structure accommodated for intratumor correlation via random intercepts. Model assumptions (Gaussian random effects and model error) were assessed via a graphical analysis of the model residuals. The synergistic effect between the two treatments was assessed via inference about the three-way interaction between time, capmatinib, and radiation.

To assess differences in longitudinal tumor growth between treatment groups, generalized linear mixed models (a Gaussian family with a log-link function) were fit to the data using the “lme4” package in R (V 4.2.3). The fixed-effect portion of the models was parameterized such that all main and interactive effects between treatment groups (vehicle control, capmatinib, radiation, and capmatinib plus radiation) and time were estimated. The random-effect structure accommodated for intratumor correlation via random intercepts for all models; random slopes were used only in the UW-lung-21model as they were found not to benefit model fit for the EBC-1 and BM06-16 data. Due to high rates of missingness (group-level sacrifice in control group), the UW-lung-21model was only fit to experimental data up to day 25. Model assumptions (Gaussian random effects and model error) were assessed via a graphical analysis of the model residuals. The synergistic effect between the two treatments was assessed via inference about the three-way interaction between time, capmatinib, and radiation.

## Results

A panel of NSCLC lines were assessed for expression of MET and phospho-MET under basal, non-stimulated conditions. The MET exon 14-mutated UW-lung-21and MET-amplified EBC-1 demonstrated higher expression of MET and phospho-MET (Fig. 1A) consistent with the known activity of these alterations to cause constitutive activation of the receptor. Response to capmatinib was evaluated using a cell proliferation assay. Both UW-lung-21and EBC-1 were inhibited by capmatinib in a dose-response manner whereas the non-MET-altered cell lines did not (Fig. 1B). The IC50 value for UW-lung-21and EBC-1 was found to be 21 nM and 2 nM, respectively, like previously reported for other MET altered NSCLC cancer cell lines(32). We then investigated the effects of capmatinib and radiotherapy on clonogenic survival. When combined with a single dose of radiation, capmatinib administered at 10 nM 1 h before radiation resulted in a significant decrease in clonogenic survival in MET altered lines UW-lung-21and EBC-1 (*P* < .01) but had no effect in the MET-wildtype line A549 or the MET-wildtype human lung epithelial line Beas-2B (*P* > 0.15) (Fig. 1C). The DEF at a surviving fraction of 0.1 for UW-lung-21and EBC-1cells exposed to 10 nM capmatinib was 1.12 and 1.26, respectively.

**Fig. 1.**
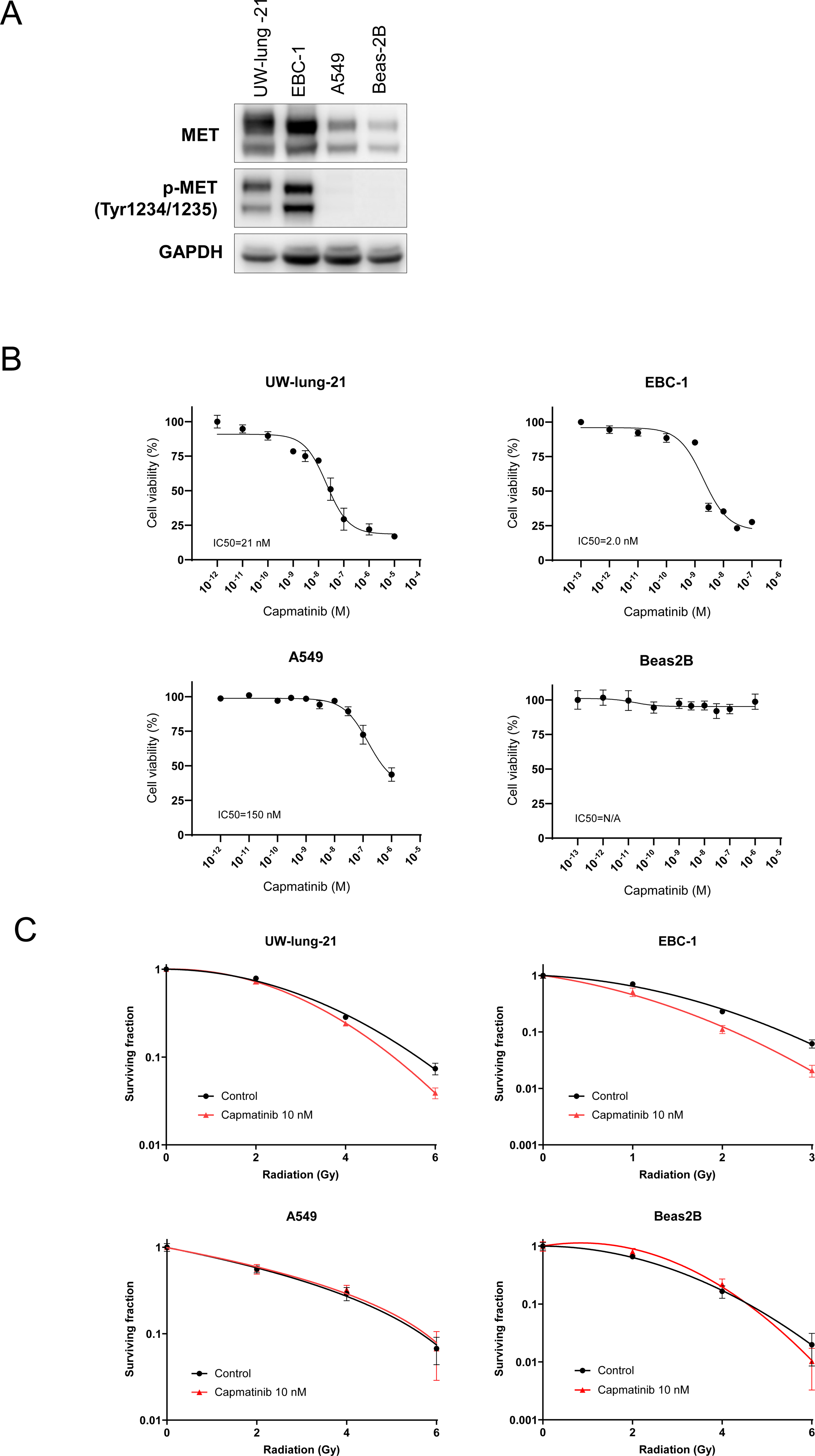
MET-altered non-small cell lung cancer (NSCLC) cell lines are responsive to capmatinib and demonstrate radiosensitization. (A) Basal levels of MET and phospho-MET were determined by western blot in a panel of NSCLC lines; UW-lung-21(MET exon 14), EBC-1 (MET amplified), A549 (MET wildtype), and Beas-2B (non-tumorigenic lung epithelial). (B) Sensitivity to capmatinib measured by proliferation assay demonstrates that MET-altered lines UW-lung-21and EBC-1 were sensitive to the MET inhibitor, while A549 and Beas-2B were significantly less sensitive to the drug. (C) *In vitro* radiosensitizing effect of capmatinib on UW- lung-21(DEF of 1.12) and EBC-1 (DEF of 1.26). Cells were treated with 10 nM capmatinib for 1 h before irradiation and maintained in the medium after irradiation. Colony-forming efficiency was determined 14 to 21 days later, and survival curves were generated. Points, mean; Bars, SEM (n = 6).

Western blot analysis was conducted to evaluate the effects of capmatinib on the activation of signaling pathways involving MET, AKT, MAPK/ERK, S6 and SAT3, both alone and in combination with radiation (Fig. 2). The results showed that capmatinib effectively inhibited phosphorylation of MET, MAPK/ERK, S6 and STAT3 in response to radiation in UW-lung-21and EBC-1 NSCLC cell lines. Each of these pathways can be activated by radiation and are involved in cell survival, proliferation, and migration. There was no response to capmatinb in MET wildtype A549 cells.

**Fig. 2.**
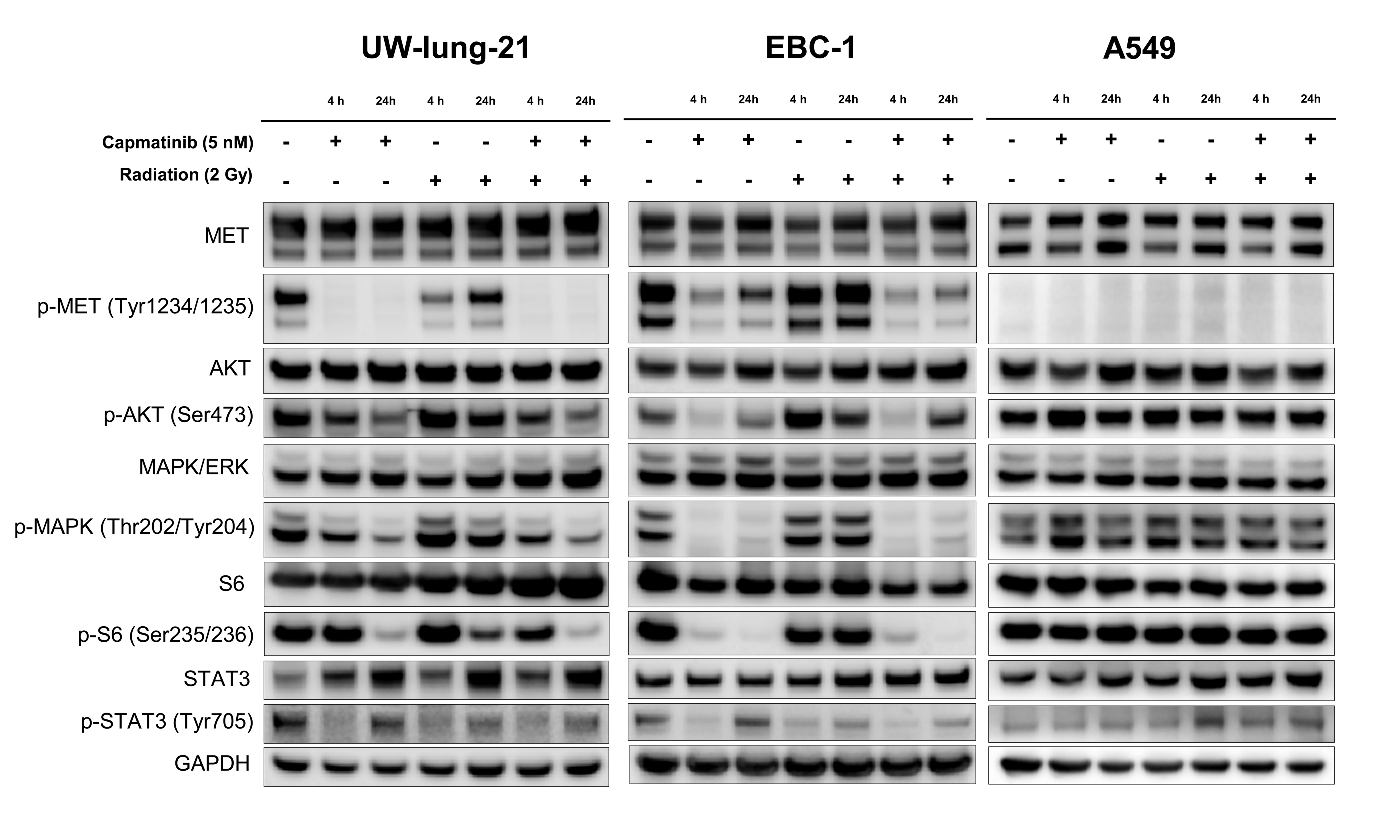
Capmatinib inhibits MET, MAPK/ERK, and S6 activation in combination with radiation. Western blot analysis was used to assess MET, p-MET (Tyr1234/1235), AKT, p-AKT (Ser473), MAPK/ERK, p-MAPK/ERK (Thr202/Tyr204), S6, p-S6 (Ser235/236), STAT3, and p-STAT2 (Tyr705) in MET-altered cell lines UW-lung-21 and EBC-1. Cells were pretreated with vehicle (DMSO) or capmatinib (5 nM) for 1 h and collected after exposure to either no radiation or radiation (2 Gy) at the indicated time points.

To determine whether capmatinib enhances radiation-induced DNA damage, we assessed the expression of γH2AX expression in the *in vitro* setting. γH2AX foci expression has been established as a sensitive indicator of DNA double-strand breaks and the dispersion of these foci has been shown to correspond to double-strand break (DSB) repair in cells exposed to irradiation. Treatment with capmatinib (10 nM) alone significantly increased γH2AX 1 h post radiation in both UW-lung-21and EBC-1 cells (*P* < .01). Capmatinib plus irradiation in UW-lung-21and EBC-1 cells resulted in a significant increase in the number of γH2AX foci at 24 h (*P* < .01) compared to cells treated with irradiation alone (Fig. 4 and Supplemental Fig 1). These findings suggest that the capmatinib-induced radiosensitization is due to the inhibition of DNA DSB repair. There was no difference in γH2AX expression in A549 cells.

γH2AX expression at 24 hours after irradiation has been shown to correlate with radiosensitivity(33), but it reflects a chromatin level response. Neutral comet assay was therefore performed to confirm capmatinib and radiation-induced DSBs. UW-lung-21and EBC1 cells, were treated with capmatinib (10 nM), radiation (4 Gy) or capmatinib (10nM) and radiation (4 Gy) collected 4 and 24 hours later. As shown in Figure 3, in both UW-lung-21(p=0.001) and EBC-1 (P<0.0001) cells exposed to capmatinib plus radiation there was a significant increase in the amount of DNA damage remaining compared with cells receiving only radiation at 24 hours. Exposure of cells to capmatinib alone had no effect on DNA damage. Thus, consistent with the γH2AX data, these results indicate that capmatinib inhibits the repair of radiation- induced DSBs.

**Fig. 3.**
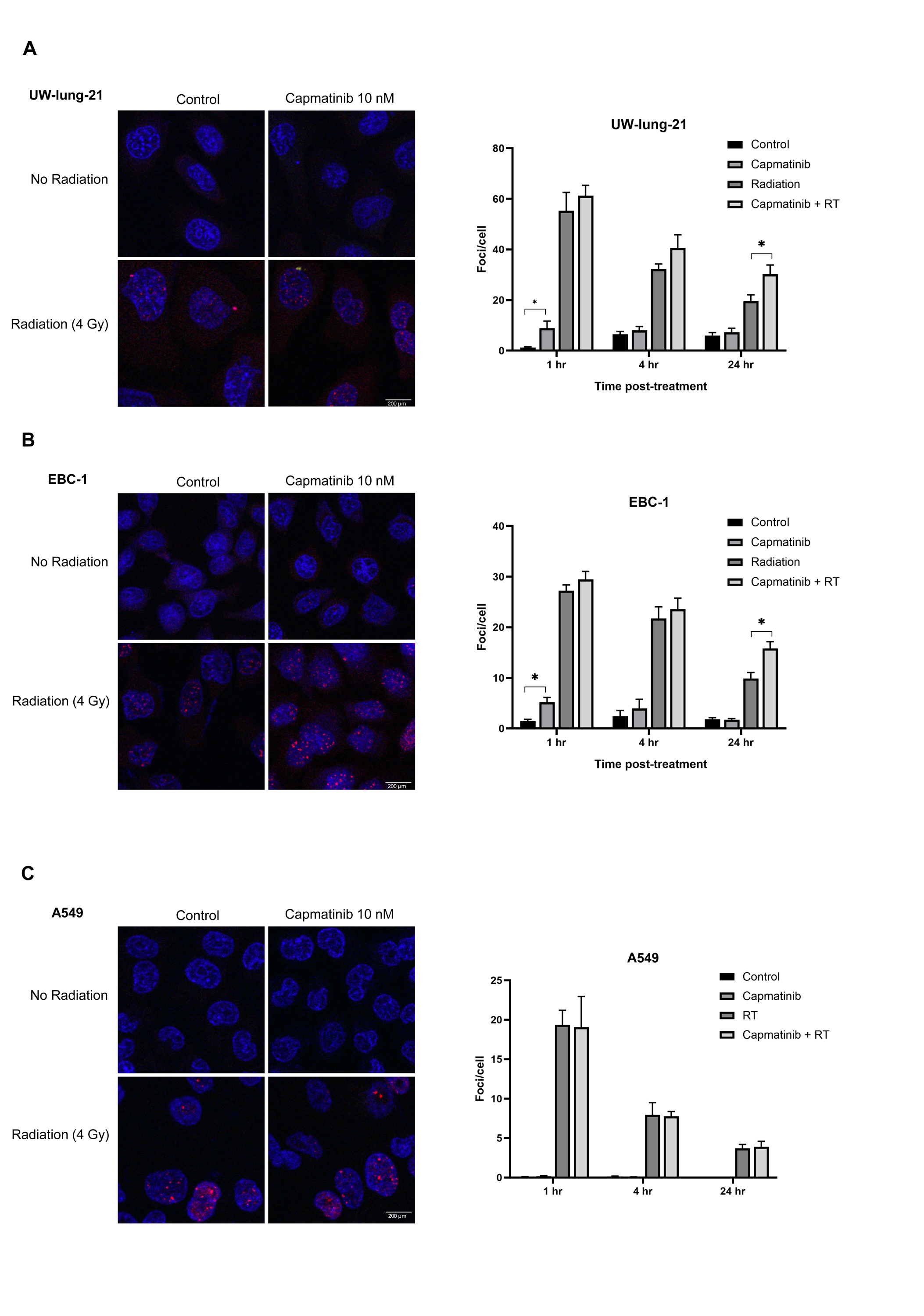
Capmatinib prolongs γH2AX foci formation induced by radiation indicating the inhibition of DNA repair. Combination of capmatinib and radiation resulted in a significant increase in the number of γH2AX foci compared to either treatment alone, indicating inhibition of DNA repair. Representative images of γH2AX foci are shown. The scale bar represents 200 µm, and all images are to the same scale. Graphs display the mean number of γh2AX foci per cell in (A) UW-lung-21and (B) EBC-1 (C) A549 cells treated with either capmatinib (10 nM), radiation (4 Gy), or the combination of both, at different time points post-treatment. The data represents the average of three independent experiments, and error bars represent standard error of the mean (SEM). *, *P* < .01, according to t test (radiation vs. capmatinib plus radiation).

To investigate whether apoptosis is involved in capmatinib-induced radiosensitization, we assessed Annexin V-FITC in UW-lung-21and EBC-1 cells 48 h following treatment with capmatinib (10 nM), irradiation (4 Gy), or both capmatinib and radiation (. As expected for a solid tumor cell line, radiation alone induced minimal apoptotic cell death. While apoptosis was increased with capmatinib plus radiation, compared to capmatinib alone or radiation alone, the overall small percentages indicate that apoptosis is not the main driver of capmatinib plus radiation-induced cell death (supplemental Fig 2).

Given our *in vitro* findings, we examined the ability of capmatinib to enhance tumor cell radiosensitivity in an *in vivo* tumor growth delay setting using three xenograft models: MET exon 14 PDX (UW-lung-21), a MET-amplified cell line xenograft (EBC-1), and in a MET-amplified PDX (BM016-16). Mice bearing subcutaneous NSCLC tumors were randomized into four groups: vehicle, capmatinib, radiation, or capmatinib plus radiation. Capmatinib (20 mg/kg) was given by oral gavage 1 h before local tumor irradiation (2 Gy x 5 daily fractions) for 5 days. The combination of capmatinib and radiation delayed the growth of tumors in all three xenografts compared with vehicle control, drug alone, or radiation alone (Fig. 5A).

**Fig 4:**
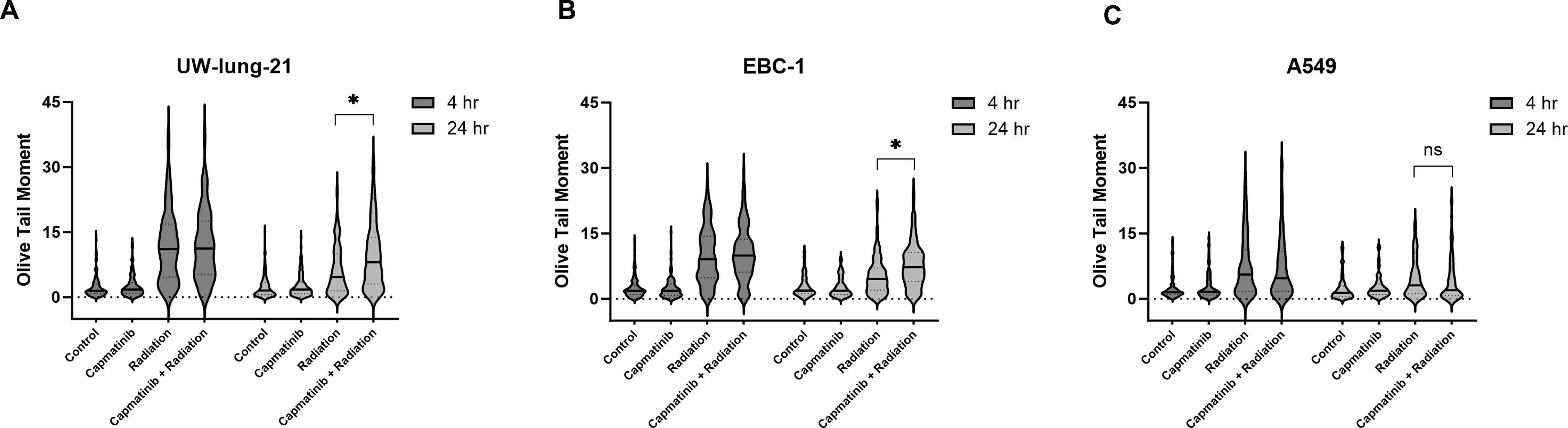
Capmatinb enhances DNA damage induced by radiation as measured by comet assay. The quantification of DSB was performed by using comet assay and measuring the Olive Tail Moment at 4 hrs and 24 hrs post treatment in (A) UW-lung-21, (B) EBC-1 and (C) A549 cells. Combination of capmatinib (10 nM) and radiation (4 Gy) resulted in a significant increase in DNA double strand breaks (DSB) measured by comet assay at 24 hours in EBC1 and UW-lung-21compared to radiation alone. There was no difference in DSB in A549 cells at 24 hrs. The data shown as Violin plots represents the average of three independent experiments, ***, *P* < .01, according to t test (radiation vs. capmatinib plus radiation).

**Fig. 5.**
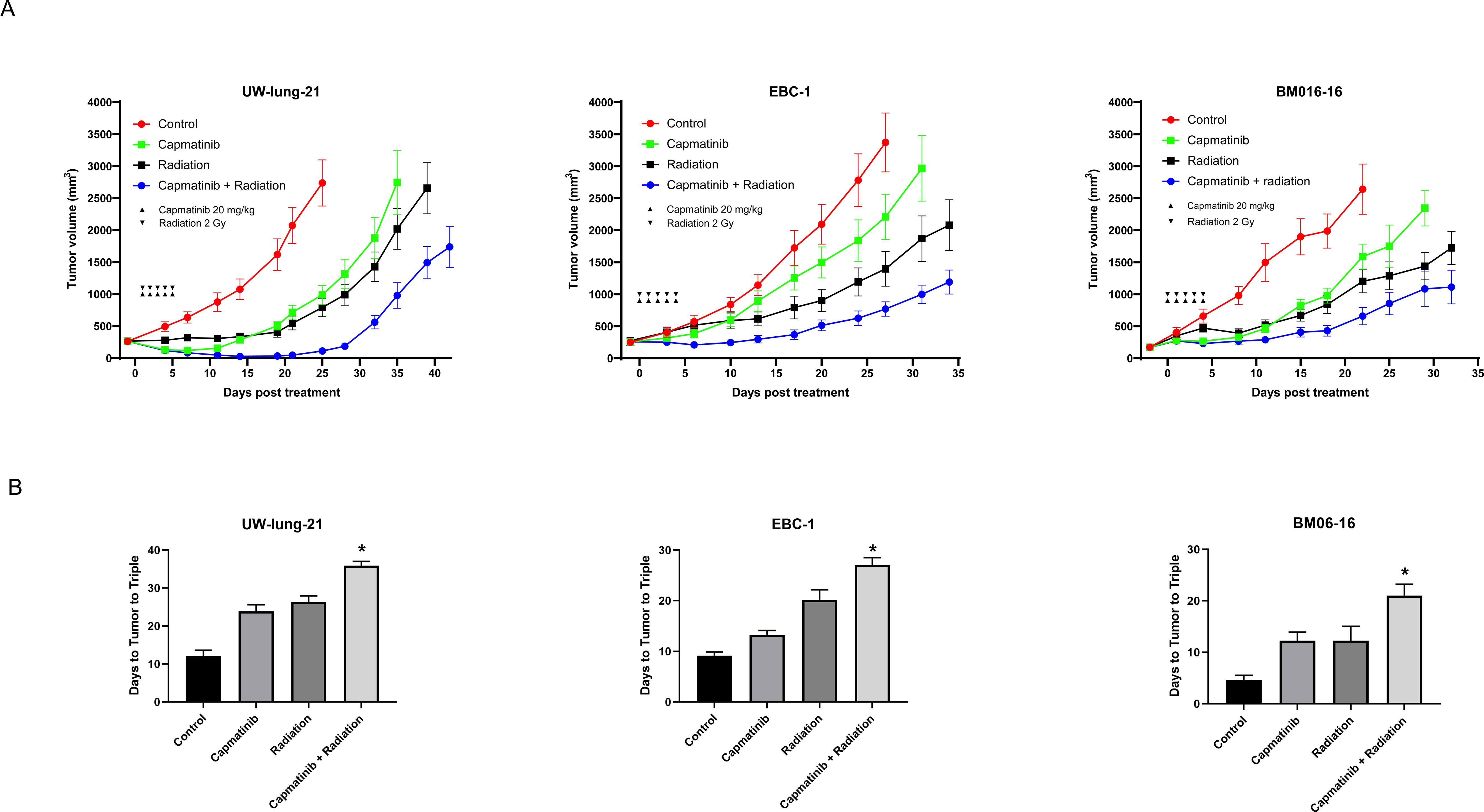
Capmatinib enhances radiation-induced xenograft tumor growth delay. (A) Tumor growth delay curves of UW-lung-21, EBC-1, and BM016-16 subcutaneous flank xenografts treated with vehicle, capmatinib (20 mg/kg), radiation (2 Gy x 5), or capmatinib (20 mg/kg) plus radiation (2 Gy x 5). Capmatinib was administered for 5 days 1 h before each radiation fraction. **(B)** Mean time in days from start of treatment until time tumor volumes tripled. Points, mean tumor volume in mice after treatment; bars, SEM. * *P* < .01 for comparison of combination group to control, capmatinib, and radiation group for all three animal studies.

The mean time for UW-lung-21PDX tumors to triple in size from start of treatment was 12.1 days for vehicle-treated mice, 23.9 days for capmatinib-treated mice, and 26.4 days for irradiated mice. In mice that received capmatinib plus irradiation, the time for tumors to triple increased to 35.9 days, which was statistically significant compared with the other groups (*P* < .001; Fig. 5B). The absolute growth delay (time in days for tumors in treated mice to triple minus the time in days for tumors to reach the same size in vehicle-treated mice) was 11.8 days for capmatinib alone and 14.3 days for irradiation alone, whereas the tumor growth delay induced by the capmatinib plus irradiation treatment was 23.8 days. For the UW-lung-21experiment, the estimated slope of the tumor growth curve for the capmatinib group was found not to be significantly different from that of the control group (*P* > .05). However, the slopes for the radiation group and the combination group were significantly lower compared to control (*P* < .01 and *P* < .001, respectively). There was evidence of a positive synergistic effect between capmatinib and radiation (*P* < .001).

For EBC-1 tumor xenografts, mean time to triple was 9.5 days for vehicle-treated mice; 12.8 days for capmatinib-treated mice; and 20.2 days for irradiated mice. In mice that received the capmatinib plus irradiation, the time for tumors to triple was 26.7 days, which was statistically significant compared with the other groups (*P* < .01; Fig. 5B). The absolute growth delay was 3.3 days for capmatinib alone, 10.7 days for irradiation alone and 17.2 days for capmatinib plus irradiation. The slope of the estimated tumor growth curve for the capmatinib group was significantly lower than that of the control group (*P* < .001). The radiation group and combination group also had growth slopes significantly less than the control group (*P* < .0001 for both comparisons), but there was no evidence of a positive synergistic effect (*P* > .05).

For the BM06-16 PDX, mean time to triple was 4.7 days for vehicle-treated mice; 10.3 days for capmatinib-treated mice; and 12.3 days for irradiated mice. In mice that received the capmatinib plus irradiation, the time for tumors to triple was 19.0 days, which was statistically significant compared with the other groups (p < .03; Fig. 5B). The absolute growth delay was 5.6 days for capmatinib alone, 7.6 days for irradiation alone, and 14.4 days for capmatinib plus irradiation. The estimated tumor growth curves for the radiation group and the combination group were both found to be significantly lower than that of the control group (*P* < .0001 and *P* < .01, respectively); the slope of the capmatinib treatment group was smaller than the control group, but this effect was not found to be statistically significant (*P* > .05), however, there was no significant synergistic effect was observed (*P* > .05).

IHC evaluated 4 h after a single treatment of capmatinib, radiation, or the combination on UW-lung-21, EBC-1, and BM06-16 tumors demonstrated inhibition of phospho-MET (Tyr1234/1235) and phospho-S6 (Ser235/236) with capmatinib alone or capmatinib plus radiation compared with control vehicle and radiation alone (Fig. 6A). In addition, Ki-67, a marker of cell proliferation, was significantly reduced in the combination group compared with the radiation alone and the other groups in UW-lung-21and EBC-1 when measured after 5 days of treatment (*P* < .05, Fig. 6B).

**Fig. 6.**
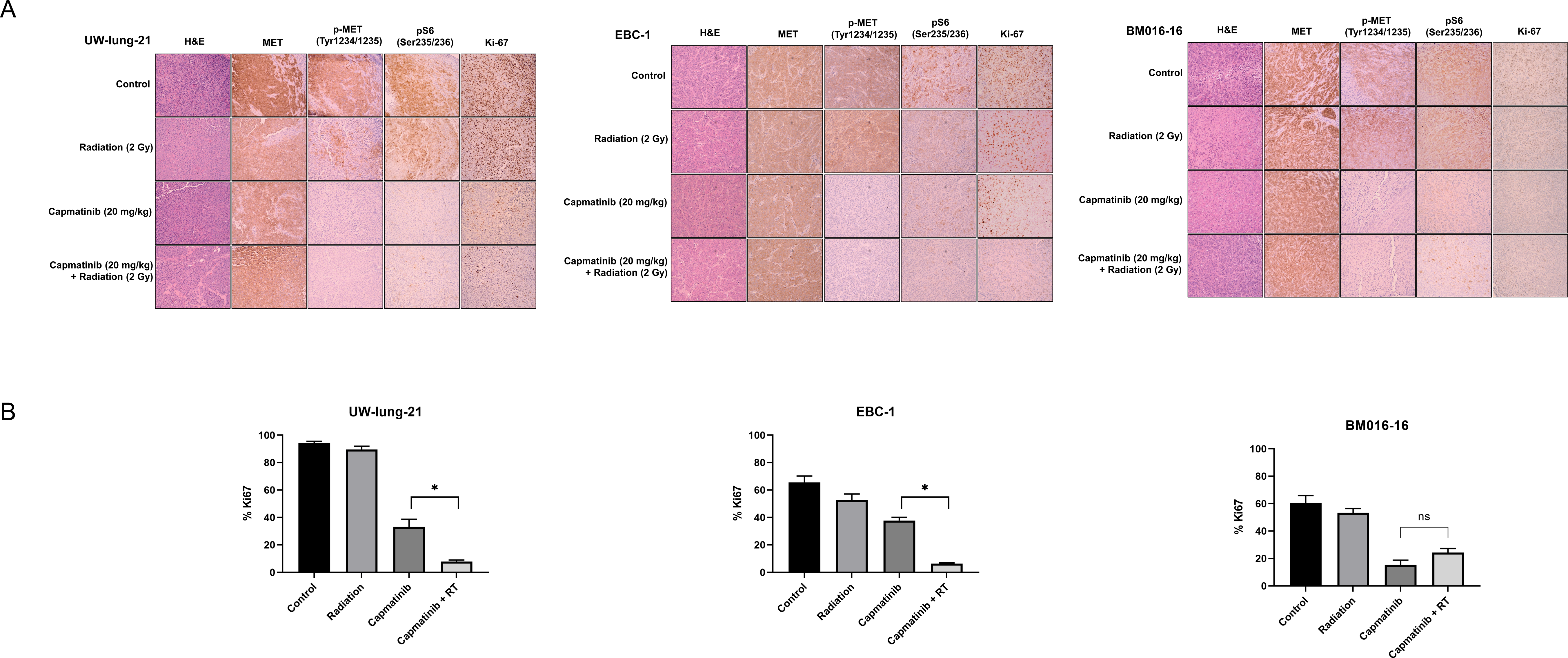
Capmatinib inhibits xenograft tumor proliferation. (A) Immunohistochemistry analysis of UW-lung-21, EBC-1, and BM016-16 tumors showing H&E, MET, p-MET (Tyr1234/1235), and p-S6 (Ser235/236) and Ki-67 expression. **(B)** Quantification of Ki-67. Tumors were harvested at 4 h after the last radiation dose 5 days following control, capmatinib (20 mg/kg), radiation (2 Gy), and capmatinib plus radiation treatment. Columns, mean; bars, SEM. * *P* < .01.

## Discussion

Given the results of GEOMETRY mono-1 study(8), capmatinib is now used as front-line therapy in patients with metastatic MET exon 14 or high MET-amplified NSCLC. Overall response rates range from 41-68% in patients with MET exon 14 skipping mutations and 29-40% in patients with a MET gene copy number of at least 10(8). While two previous small studies evaluating non-clinical MET inhibitors in NSCLC have suggested potential radiosensitization, it is unclear whether capmatinib induced the same radiation enhancement effect, which could have potential implications when combining capmatinib and radiation in the clinic.

In this study, we provide preclinical evidence of radiosensitization with the selective MET inhibitor capmatinib in both MET exon 14-mutated and MET-amplified NSCLC models. The combination of capmatinib with radiation resulted in reduced DNA DSB repair and enhanced tumor growth delay in our MET-altered PDX models. Our results are important because we evaluated a clinically relevant MET inhibitor, are the first to demonstrate radiation enhancement in a MET exon 14-mutated model, and we validated our results with MET exon 14 and MET-amplified PDX studies.

Besides capmatinib, tepotinib and crizotinib are also approved for use in the United States, with savolitinib being approved in China for patients with NSCLC harboring MET exon 14 skipping alterations. Crizotinib is an unselective MET inhibitor and two studies have failed to show radiosensitization with crizotinib(34, 35). Others have shown that tepotinib has preclinical synergism with radiation in MET expressing head and neck cancers(25) and glioma models(36). We have previously demonstrated that salovitinib enhances the effect of radiation in our MET exon 14 PDX model(26) further supporting radiosensitization with selective MET inhibition.

MET is an attractive target for radiosensitization given radiation upregulates MET transcription and overexpression through modulation of ATM and NF-κB(37), leading to receptor phosphorylation and signal transduction pathway activation (e.g., RAS/RAF/MAPK, PI3K/AKT, and STAT), resulting in enhancement of DNA repair(38–41). In addition, MET expression correlates with increased hypoxia, which is a powerful factor in resistance to ionizing radiation therapy.(42) The proposed mechanisms of MET-induced radiosensitization include inhibition of homologous repair of double-strand DNA breaks through sustained activation of γ-H2AX(16, 18, 19) and reduced levels of Rad51(41) as well as reduced phosphorylation of ATR and CHK1 abrogating an associated DNA damage–induced S phase arrest(19). Our data support the impairment of DNA repair as a main mechanism of capmatinib induced radiation sensitization. Further work will be needed to fully elucidate and exploit this process.

## Conclusion

In summary, we have demonstrated both *in vitro* and *in vivo* radiation enhancement by combining the MET inhibitor capmatinib with clinically relevant doses of radiotherapy in MET exon 14 and MET-amplified NSCLC models. This combination holds translational implications, given the well-established use of radiotherapy in the treatment of NSCLC and the use of capmatinib in patients with MET alterations. In locally advanced MET altered NSCLC, MET inhibitors could be given concurrently with radiation as a radiosensitizer and then as adjuvant therapy to reduce the risk of recurrence. In the metastatic setting ablative radiation could be used to control macroscopic disease while MET TKIs could be used to control disseminated microscopic disease. Overall, these data provide the rationale for continuation of studies combining radiotherapy with MET inhibition. Ultimately, clinical trials will provide important information on the safety and effectiveness of this combination and will help guide future treatment decisions for patients with NSCLC.

## Appendix A. Supplementary data

Supplementary data to this article can be found online.

## Data Availability Statement for this Work

Research data are stored in an institutional repository and will be shared upon reasonable request to the corresponding author.

## Acknowledgements

The authors would like to thank James P. Zacny, PhD for manuscript preparation and formatting assistance.

