## Supplemental Data for "MET Inhibitor Capmatinib Radiosensitizes MET Exon 14-Mutated and MET-Amplified Non-Small Cell Lung Cancer"

**Supplemental Table 1: short tandem repeat profiling of cell lines**

| | UW-lung 21 | | | EBC-1 | Beas-2B | A549 |
|---|---|---|---|---|---|---|
| | P0* | Genetica: 5/3/21 | UW TRIP: 5/24/22 | UW TRIP Lab: 2/8/22 | Genetica: 8/20/19 | UW TRIP Lab: 5/3/23 |
| **FGA** | 21,21.2 | 20,21,21.2 | 20,21,21.2 | 22,22 | 20, 24 | 23,23 |
| **TPOX** | 8,9 | 8,9 | 8,9 | 8,8 | 6, 11 | 8,11 |
| **D8S1179** | 12,16 | 11,12,13,16 | 11,12,13,16 | 14,14 | 13, 15 | 13,14 |
| **vWA** | 14,16 | 14, 16 | 14,16 | 16,17 | 17, 18 | 14,14 |
| **Amelogenin** | X, Y | X, X | X, X | X,X | X, Y | X, X |
| **Penta_D** | 12,14 | 10,12,13,14 | 10,12,13,14 | 13,13 | 2.2, 13 | 9,9 |
| **CSF1PO** | 10,12 | 10 | 10,10 | 10,10 | 9, 12 | 10,13 |
| **D16S539** | 12,12 | 12,13 | 12,13 | 9,9 | 12 | 11,12 |
| **D7S820** | 10,11 | 10,11,12 | 10,11,12 | 10,11 | 10, 13 | 8,11 |
| **D13S317** | 10,10 | 10,11 | 10,11 | 12,12 | 13 | 11,11 |
| **D5S818** | 11,12 | 11,12 | 11,12 | 11,11 | 12, 13 | 11,11 |
| **Penta_E** | 8,15 | 7,8,15,16 | 7,15,16 | 16,16 | 5, 8 | 7,11 |
| **D18S51** | 15,21 | 17,21 | 17,21 | 12,17 | 18, 19 | 14,17 |
| **D21S11** | 29,30 | 28,29,30,33.2 | 28,29,30,33.2 | 29,30 | 28, 30 | 29,29 |
| **TH01** | 9,9.3 | 9,9.3 | 8,9,9.3 | 7,7 | 7, 9.3 | 8,9.3 |
| **D3S1358** | 16,16 | 16,17,18 | 16,17,18 | 15,15 | 15, 17 | 16,16 |
| Passage # | P0 | P5 | P15 | P7 | P14 | P17 |

***Baschnagel et. al., Sci. Rep. 2021**

**Supplemental Table 2: List of cell lines used, sources, and culture conditions.**

| Cell line | Source | Culture condition |
|---|---|---|
| EBC-1 | JCRB (Japanese Collection of Research Bioresources) Cell Bank Catalog #: JCRB0820 | EMEM with 200 mM L-glutamine, 10% FBS, penicillin (100 units/mL), streptomycin (100 mg/mL) |
| UW-lung 21 | Established in Baschnagel and Kimple lab, University of Wisconsin | ATCC modified RPMI-1640 (catalog #: 30-2001), 10% FBS, penicillin (100 units/mL), streptomycin (100 mg/mL) |
| A549 | ATCC (Catalog #: CCL-185) | DMEM/F12, 10% FBS, penicillin (100 units/mL), streptomycin (100 mg/mL) |
| Beas-2B | ATCC (Catalog #: CRL-9609) | RPMI, 10% FBS, penicillin (100 units/mL), streptomycin (100 mg/mL) |

**Supplemental Table 3: Antibodies**

| Antibody | Source | Catalog # | Company | Dilution | Detection method |
|---|---|---|---|---|---|
| MET | Rabbit | CST 8198 | Cell Signaling Technology | 1:1000 | Western Blot |
| phospho-Met (Tyr1234/1235) | Rabbit | CST 3077 | Cell Signaling Technology | 1:500 | |
| AKT | Rabbit | CST 2920 | Cell Signaling Technology | 1:1000 | |
| phospho-AKT (Ser473) | Rabbit | CST 4060 | Cell Signaling Technology | 1:1000 | |
| p44/42 MAPK (Erk1/2) | Rabbit | CST 4696 | Cell Signaling Technology | 1:1000 | |
| phospho-p44/42 MAPK (Erk1/2) (Thr202/Try204) | Rabbit | CST 4370 | Cell Signaling Technology | 1:500 | |
| S6 | Rabbit | CST 2217 | Cell Signaling Technology | 1:1000 | |
| phospho-S6 (Ser235/236) | Rabbit | CST 4858 | Cell Signaling Technology | 1:1000 | |
| STAT3 (Tyr705) | Mouse | SC-8019 | Santa Cruz Biotechnology | 1:1000 | |
| phospho-STAT3 | Mouse | SC-8059 | Santa Cruz Biotechnology | 1:500 | |
| GAPDH | Rabbit | CST5174 | Cell Signaling Technology | 1:1000 | |
| MET | Rabbit | CST 8198 | Cell Signaling Technology | 1:300 | Immunohistochemistry |
| phospho-Met (Tyr1234/1235) | Rabbit | CST 3077 | Cell Signaling Technology | 1:300 | |
| phospho-S6 (Ser235/236) | Rabbit | CST 4858 | Cell Signaling Technology | 1:400 | |
| Ki-67 | Rabbit | CST 9027 | Cell Signaling Technology | 1:400 | |
| phospho-Histone H2AX | Rabbit | CST 9718 | Cell Signaling Technology | 1:400 | Immunofluorescence |
| Alexa Fluor 546 | Rabbit | A-11035 | Invitrogen | 1:500 | |

**Supplemental Figure 1: UW-lung-21 γH2AX Images**

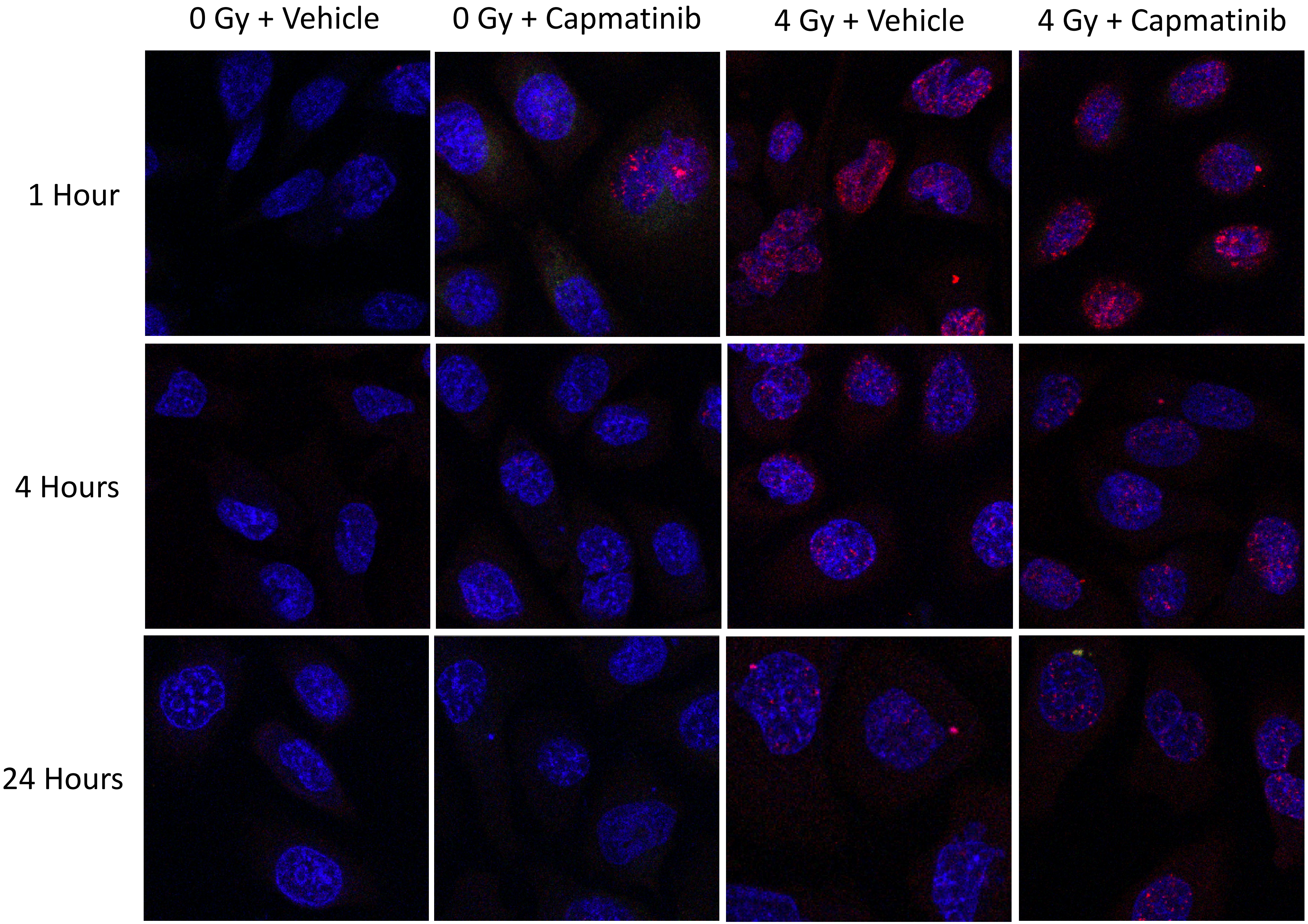

**Supplemental Figure 1: EBC1 γH2AX Images**

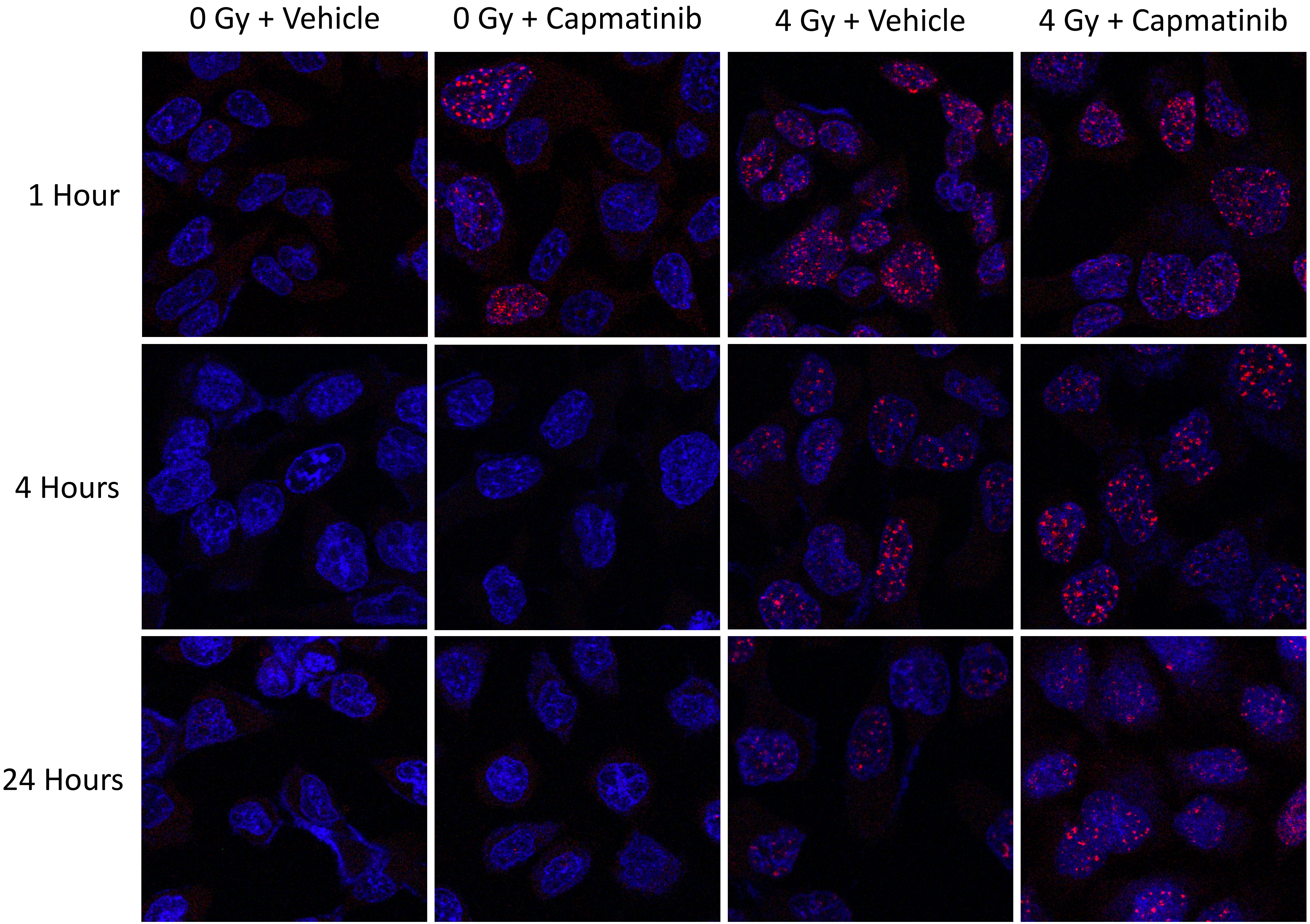

**Supplemental Figure 1: A549 γH2AX Images**

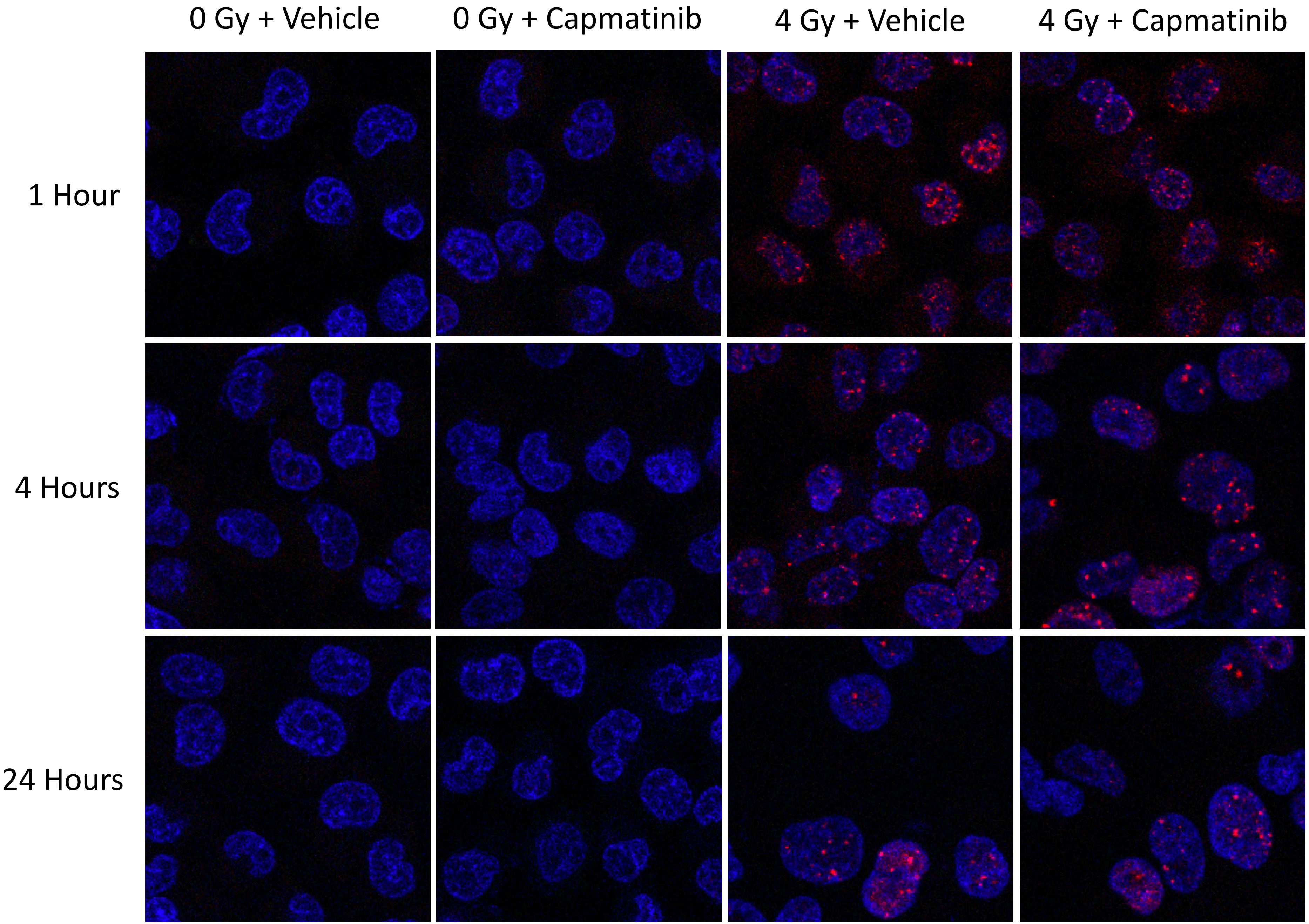

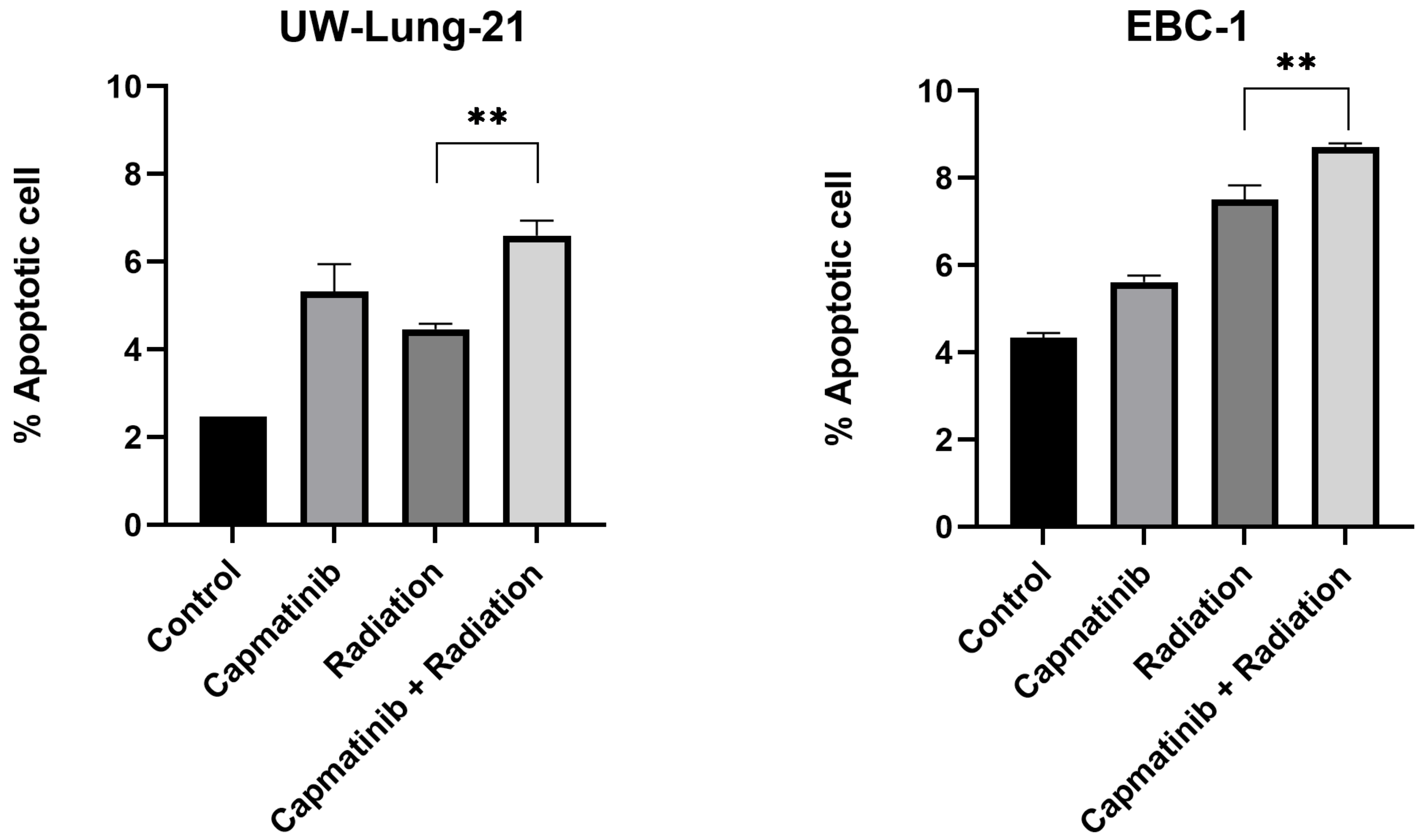


**Supplemental Figure 2. Capmatinib increases apoptosis in combination with radiation.** Cells were treated with vehicle (DMSO) or capmatinib (10 nM) 1 h prior to 4 Gy radiation and analyzed at 48 h time point. The total percentage of apoptotic cells combining early and late apoptosis was measured by flow cytometry using Annexin V and FITC staining kit. Columns, mean; bars, SEM (n=3). *, $P < .01$, according to t test (radiation vs. capmatinib plus radiation).
